# Rhythmic entrainment source separation: Optimizing analyses of neural responses to rhythmic sensory stimulation

**DOI:** 10.1101/070862

**Authors:** Michael X Cohen, Rasa Gulbinaite

## Abstract

The so-called steady-state evoked potentials (SSEPs) are rhythmic brain responses to rhythmic sensory stimulation, and are often used to study perceptual and attentional processes. We present a data analysis method for maximizing the signal-to-noise ratio of the narrow-band steady-state response in the frequency and time-frequency domains. The method, termed rhythmic entrainment source separation (RESS), is based on denoising source separation approaches that take advantage of the simultaneous but differential projection of neural activity to many non-invasively placed electrodes or sensors. Our approach is a combination and extension of existing multivariate source separation methods. We demonstrate that RESS performs well on both simulated and empirical data, and outperforms conventional SSEP analysis methods based on selecting electrodes with the strongest SSEP response. We also discuss the potential confound of overfitting—whereby the filter captures noise in absence of a signal. Matlab scripts are available to replicate and extend our simulations and methods. We conclude with some practical advice for optimizing SSEP data analyses and interpreting the results.

---

The steady-state evoked potentials (SSEPs) are rhythmic neural responses to rhythmic sensory stimulation. It has many applications in vision, cognitive, and clinical neuroscience research, as well as brain computer interfaces (for reviews, see Norcia et al., 2015; Vialatte et al., 2010). Our focus here concerns a novel SSEP data analysis method that is aimed to increase utility of SSEPs in vision and cognitive neuroscience research.

The traditional data analysis approach in SSEP research involves examining power spectra or narrow-band activity from a single or several EEG electrodes (or MEG sensors) that show maximal response to rhythmic stimulation (“best-electrode” approach; e.g. Fuchs et al., 2008). This seemingly simple and straightforward approach has several data analysis issues, which we detail below. When exploring various analysis procedures, we found that none is particularly well-suited to resolve these issues. However, a combination of existing temporal and spatial filtering methods outperforms “best-electrode(s)” approach both in simulations and in real data. The purpose of this paper is to present this SSEP data analysis approach, termed rhythmic entrainment source separation (RESS), and provide specific recommendations for SSEP experiment design and data analysis. RESS is a combination of several existing methods, principally including those from Nikulin et al. (Nikulin et al., 2011), Chevinge et al. (de Cheveigné and Arzounian, 2015; de Cheveigné and Parra, 2014), as well as others (Dmochowski et al., 2015; Särelä and Valpola, 2005).

The following list summarizes data analysis challenges in SSEP research, and ways that our RESS procedure resolves those challenges. Many points are focused on the steady state *visual* evoked potentials (SSVEPs), but are applicable to SSEPs in other sensory modalities as well.

1. *Electrode selection.* Although most experiments involve recording brain activity from many dozens of electrodes, a single or several electrodes are generally selected for SSEP data analyses (Andersen et al., 2011; Rossion et al., 2012). But the “best” electrode may be difficult to select, may require some subjective judgment, and differs across stimulation frequencies and across individuals. To address this issue, we construct spatial filters that take weighted combinations of all electrodes to produce a single time series (“RESS components”).
2. *Small effects.* High signal-to-noise SSVEPs are acquired using low flicker frequencies (<30 Hz), and when using large, high-luminance stimuli that flicker for several to tens of seconds (Kashiwase et al., 2012; Painter et al., 2014; Srinivasan et al., 2006). However, cognitive paradigms often require smaller stimuli and short presentation times (Gulbinaite et al., 2014; Peterson et al., 2014). The resulting SSVEPs might be difficult to isolate from ongoing brain activity in the same frequency ranges. We show that generalized eigendecomposition in combination with careful selection of a reference covariance matrix can boost signal-to-noise ratio of SSEPs and thus facilitate the use of SSVEPs in cognitive paradigms.
3. *Separating endogenous oscillations from the sensory-entrainment response.* When the flicker frequency overlaps with frequencies of endogenous brain oscillations (e.g., in the range of 2-40 Hz), the SSEPs can be difficult to disentangle from the endogenous brain activity. Although for some experiments it might be suboptimal to have flicker in the range of brain oscillations, other studies have capitalized on this to investigate whether sensory flicker can interfere with ongoing neural oscillations (Spaak et al., 2014). We show that RESS can be used to subtract SSEP-related activity from the data while leaving endogenous activity relatively intact (or at least, more intact compared to a notch filter).
4. *Time series analysis of the time-course of the SSEP.* If the amplitude of the SSEP indexes attention differences across conditions, it is reasonable to expect that time-varying fluctuations in SSEP amplitude might reflect time-varying fluctuations in attention, which can be extracted using narrow band-pass filtering (Andersen and Müller, 2010). However, very-narrow-band filtering obliterates fine temporal dynamics, because temporal non-stationarities are represented by side-lobes of the frequency spectrum, and not by the peak at the flicker frequency. We show that by applying a spatial filter to non-temporally filtered or weakly filtered data allows accurate reconstruction of the time course of SSEP amplitude fluctuations.
5. *Separating SSEPs for multiple simultaneous frequencies.* Different flicker frequencies are often presented simultaneously in order to track attention to multiple items. Providing the spatiotemporal dynamics of the SSEPs are distinct, the RESS procedure can isolate activity from multiple simultaneous flicker frequencies, even those separated by 1 Hz.
6. *Single-trial analyses.* The RESS method is optimized for within-subject cross-trial analyses, while most SSEP analysis methods are optimized for trial-average analyses (or are sometimes strictly limited to trial-average results). Indeed, the RESS method does not require multi-trial designs. This means that the onset phase of the flicker can differ on each trial, which increases experiment and analysis flexibility.

Finally, we stress that the RESS procedure is not a “black-box toolbox” that should be applied to data without thinking. Instead, we recommend careful inspection of the data, careful selection of analysis parameters, and an understanding of how and why the method works. To this end, we provide commented Matlab code that can be used, adapted, and extended. In this sense, RESS is more suitable for vision and cognitive and neuroscience studies that involve off-line data analysis.

## Methods

### Overview of RESS

The general approach of using “guided” (as opposed to blind) source separation techniques in multivariate signals has a decades-old tradition. In this sense, the method we developed and present here is not a new technique per se, but instead, is a blend of several existing techniques that are optimized for SSEP data analysis. The procedure is outlined in Fig. 1. Matlab code to implement and modify the method, as well as to run the simulation and apply to sample EEG data, are provided online Data and scripts are currently available at mikexcohen.com/ress.zip (eventually, this zip will be moved to data.donders.ru.nl).

**Fig. 1.**
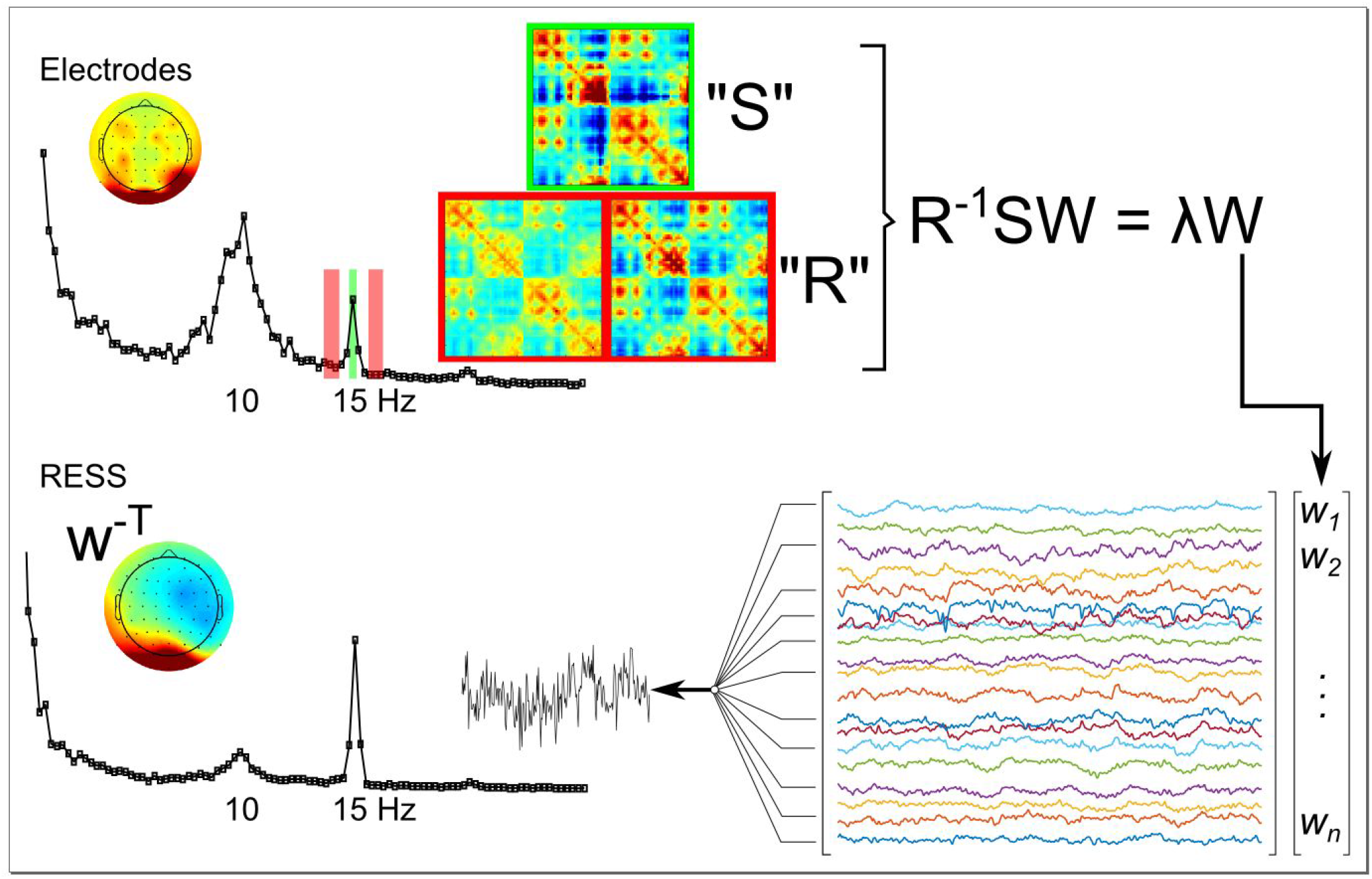
Overview of RESS procedure. Starting with the electrode data (top left), channel-to-channel covariance matrices are constructed from the data temporally filtered at SSEP frequency (15 Hz in this example), and at closely spaced neighboring frequencies (green and red covariance matrices). A generalized eigendecomposition is then computed to identify eigenvectors (“W”) that separate the signal (“S”) covariance from the average of the two flanking (“R”) covariance matrices. The eigenvector with the largest eigenvalue is taken as a spatial filter. That eigenvector is then multiplied by the raw channel data to produce a single time series, the RESS component. This time series can then be analyzed in the time or in the frequency domains, and can be visualized in topographical or anatomical maps.

The core idea is to construct linear spatial filters (RESS filters), which are then multiplied by the EEG electrode time series data to produce *RESS component* time series – a weighted combination of the electrodes – that can be analyzed instead of the data from individual electrodes. The *RESS filters* are defined by eigenvectors extracted from covariance matrices. This procedure is conceptually similar to principle components analysis (PCA), where the eigenvector corresponding to the largest eigenvalue from a covariance matrix points in the direction of maximal variance of the data. However, instead of using a single covariance matrix, we start with two covariance matrices and use generalized eigendecomposition to find a vector that maximally differentiates the two matrices. One can think of this procedure as finding a filter that maximizes a multivariate signal-to-noise ratio. Optimizing the filter for a specific application then becomes a matter of selecting the best data to use to construct the signal covariance (matrix **S)** and the “reference” covariance (matrix **R)** (we prefer the term “reference” over “noise”). For RESS, we construct **S** from data that are temporally filtered at the SSEP frequency, and we construct **R** from data that are either filtered at frequencies neighboring the SSEP frequency or are computed from the broadband signal (temporal filtering procedures are described below). Using data from different frequencies but the same time window is preferable to computing **R** from a separate baseline time period, because all cognitive, perceptual, behavioral, and other factors are held constant; the matrices differ only by their frequency content.

In theory, we want to compute the eigendecomposition of the matrix product **R**^-1^**S**, where ^-1^ is the matrix inverse. For numerical stability reasons, the better solution is to compute **SW = ΛWR,** where **W** is a matrix of eigenvectors and **Λ** is a diagonal matrix of eigenvalues. This is implemented in Matlab as `[w, L] =eig (s, R)`. The column of the **W** matrix associated with the largest eigenvalue in **Λ** is a spatial filter of size electrodes-by-1 that, when multiplied by the electrode time series, produces the largest RESS component, which is then used in data analyses. Note that unlike with PCA, RESS components are not pairwise orthogonal. This is because although **S** and **R** are both symmetric matrices, **R**^-1^**S** is generally not symmetric, and eigenvectors of a non-symmetric matrix are generally not orthogonal. This is advantageous because the orthogonality constraint is often suboptimal in neuroscience data analyses.

Because the data used to create **S** are narrow-band filtered and provide little temporal information, we take the suggestion of Cheveigne and Arzounian (2015) to apply the spatial filter to the raw data, rather than to the temporally filtered data. This maximizes temporal precision, and allows accurate reconstruction of the SSEP time course.

The major assumption of the RESS method is that the SSEP activity is spectrally and spatially stationary over time. Spectral stationarity is provided by the stationarity of the rhythmic stimulus, and thus this assumption is easily met (our filtering method is robust to the minor nonstationarities that capture the temporal fluctuations in amplitude). The spatial stationarity assumption would be violated, for example, if a flickering visual stimulus moved around on a screen and therefore elicited activity in different parts of visual cortex, with consequently different projections to the scalp. Spatial stationarity is important because RESS is a spatial filter.

Matrix regularization is sometimes applied when creating spatial filters based on eigendecomposition (Lotte and Guan, 2011), and can be helpful in situations of near-singular or high-conditioned matrices. We explored the effects of Tikhonov regularization with various parameters, and observed little effect on the results. This is likely due to the robust effect of flicker on the EEG signal. Regularization was therefore not included in any of the analyses reported here, but we leave open the possibility that regularization may be helpful in other situations, e.g., for very small SSEP effects or when using very high-density EEG arrays.

### Temporal filter

We used a frequency-domain Gaussian filter, in which the Fourier transform of the EEG signal is point-wise multiplied by a Gaussian with a specified peak and full width at half maximum (FWHM), and the inverse Fourier transform is then applied to recover the time-domain bandpass filtered signal. This procedure is also called circular convolution implemented in the frequency domain. Other types of bandpass filters could be suitable, such as FIR or IIR. We prefer a Gaussian filter because it is fast and easy to implement, has no sharp edges that would produce artifactual ripples, and is robust and easy to work with. For example, FIR filters would require additional supervision to ensure reasonable frequency responses at appropriate order parameters, and IIR filters can exhibit phase instabilities. In the Results section, we report ranges of FWHM, which are defined symmetrically around the filter frequency peak.

There is a reason to prefer a Gaussian-shaped filter and time-domain covariance over the cross-spectral density from a single Fourier coefficient. The brain's response to flicker is not perfectly stationary, nor is it perfectly sinusoidal. These non-sinusoidal and non-stationary characteristics produce a slightly wider representation in the frequency domain, and thus focusing exclusively on a single Fourier coefficient at the stimulation frequency will lead to a loss of signal. Similarly, we use the total covariance (average of each trial's covariance) rather than the covariance of the time-domain average, because the non-stationarities differ over trials resulting from fluctuations in attention, among other factors.

### Simulated data

Simulations are advantageous because the ground truth is known. On the other hand, simulations often fail to represent characteristics of real data. Our solution to this is to use an empirical EEG dataset as “realistic noise,” and add a sine wave as the simulated SSEP. The empirical data were taken from an eyes-open rest recording in an adult human volunteer (120 seconds). The SSEP was simulated as an amplitude-modulated sine wave that was added to a dipole in the brain, and then projected onto the scalp electrodes. The purpose of the amplitude modulation was to simulate time-varying fluctuations in attention. The overall goals of the simulation were to examine how the analysis parameters affect the static (power spectrum of the data) and dynamic (time-varying fluctuations of power at the flicker frequency) SSEP characteristics.

### Empirical data

We collected empirical data from one human volunteer (author RG) using a variety of experimental conditions designed to assess the utility of RESS for SSEP data analysis. Data were acquired from only one subject as a proof-of-principle demonstration of the method. The experiment involved recording EEG activity from 64 electrodes (BioSemi equipment; see biosemi.com for hardware details), sampled at 1024 Hz. Visual stimulation was delivered by a custom-built setup containing: Fixation LED light placed at the level of the eyes, and two LED arrays placed in the lower visual field with an eccentricity of 6.5 visual angle relative to the fixation LED. Auditory stimulation was delivered via headphones. There was no cognitive task required other than maintaining attention. Additional descriptions of various stimulation conditions are provided in the Results section. Offline, EEG data were high-pass filtered at .1 Hz. No other preprocessing was applied, and the raw data thus contain typical EEG artifacts such as blinks. RESS suppresses these artifacts.

### Component subtraction

Given research interests that involve studying interactions between SSEPs and the spontaneous EEG activity (Birca et al., 2006; Mast and Victor, 1991), it would be ideal to separate the two signals. In practice, a perfect separation seems unlikely. In part this is because the rhythmic stimulation interacts with ongoing endogenous oscillations and produces some entrainment of neural oscillations, particularly when the stimulation frequency matches an endogenous neural oscillation frequency (Antal and Herrmann, 2016; Notbohm et al., 2016).

Nonetheless, attempted separation of SSEP and endogenous oscillatory activity can be implemented by removing the largest RESS component(s) from the raw data. Although it is intuitive to conceptualize *subtracting* a component from data, in fact, it is implemented by *reconstructing* the data using N-1 components. The reconstruction is given by **V^-T^V^T^D,** where matrix **V** is the eigenvectors matrix **W** with the column associated with the largest eigenvalue—corresponding to the eigenvector used to obtain the RESS component—removed, ^T^ indicates the transpose, ^-T^ indicates the inverse of the transpose, and **D** is the channels-by-time (unfiltered) data matrix. (Note that if no components are removed, the above equation will simply perfectly reconstruct the data, because **W^-T^W^T^ = I.)**

### Topographical and anatomical localization

The components framework taken here allows for visualizing scalp-level topographies and putative anatomical generators associated with each RESS component. Because the term “source” is sometimes used for anatomical localization (e.g., the result of beamforming) and sometimes used for statistical isolation (e.g., blind source separation), we use the term “anatomical localization” to refer to the putative distribution of locations or possible generators in the brain. Scalp-level (topographical) visualization of the RESS filters is accomplished by computing its “activation pattern” (Haufe et al., 2014) or forward model. For a full-rank covariance matrix, the forward model is contained in the columns of **W^-T^.** For reduced-rank covariance matrices, the forward model is contained in the columns of the matrix product **SW(W^T^SW)**^-1^ (note that in the case of all full-rank matrices, this becomes **SWW^-1^S^-1^W^-T^** which simplifies to **W^-T^).** To facilitate visual interpretation of color, we force the component sign by setting the electrode with the largest forward model value to be positive.

Brain-level visualization based on 64 electrodes without precise electrode location measurements or a subject-specific head model entails some uncertainty and spatial smoothing (Fuchs et al., 2002), but still provides useful information that is more anatomically interpretable compared to topographical maps. We estimated anatomical localization of the components by adapting a procedure developed for obtaining distributed localization of independent components (Hild and Nagarajan, 2009), which mainly involves correlating the forward model of the RESS component with a leadfield matrix developed for the EEG montage. To increase sensitivity, our leadfield contained three cardinal orientations per voxel, and correlation magnitude was taken as the length of the projection of the correlation onto the dipole orientation space. The leadfield was created from a standard MRI and BEM model, using the Matlab Brainstorm toolbox (Tadel et al., 2011), which in turn utilizes algorithms developed by openmeeg (Gramfort et al., 2010). Because the components are a weighted combination of all electrodes, thresholding the anatomical distribution based on inferential statistical values makes little sense. Instead, to facilitate visual inspection, we assign color values only to voxels that have a correlation magnitude greater than the median correlation over all voxels. A brain mesh was then created in Matlab with patches being colored according to correlation magnitude intensity or to gray for subthreshold patches.

### Quantifying the static and dynamic SSEP

We use the term “static SSEP” to refer to the power spectrum of the data (extracted via the fast Fourier transform) recorded over a period of time. In this sense, “static” indicates that we extract no information about temporal dynamics. The static SSEP is computed as SNR, which we define as the ratio between power at the SSEP frequency, to power at the average of neighboring frequencies (+/- 2 Hz, excluding +/- .5 Hz around the SSEP frequency). Converting to SNR units has several advantages, including accounting for 1/f power spectral shifts, robustness to electrode montage choices, and comparability across individuals, groups, and methods (EEG, MEG, LFP). In figures we show the SNR spectrum, which is the SNR computed at each frequency.

We use the term “dynamic SSEP” to refer to time-varying fluctuations of power at the SSEP frequency. This time course is obtained by extracting the magnitude of the Hilbert transform of the narrow-band filtered time course (see also the previous section on *Temporal Filter).* For the simulated data, it is possible to test the accuracy of the reconstruction by comparing the electrode data to the simulated dipole activity. This was implemented as R^2^, where a value of 1 indicates a perfect reconstruction.

## Results

### Isolating and subtracting a simulated SSVEP

We simulated SSVEP data by generating a sine wave with smoothed random time-varying amplitude, projecting this signal to 64 scalp EEG electrodes from a dipole placed in the occipital lobe, and then summing this signal with the real EEG data taken from 120 seconds of resting state EEG recording in a human volunteer. The **S** covariance matrix was generated after applying a narrow-band-pass filter around the SSVEP frequency. The **R** covariance matrix was computed either from the average of neighboring frequencies (above and below the SSVEP frequency), or from the broadband (unfiltered) signal. The resulting spatial filter was then applied to the broadband data, from which the power spectrum was obtained via the Fourier transform.

RESS successfully isolated the SSVEP component, with near-total annihilation of activity at all other frequencies (Fig. 2A). We repeated this simulation using a variety of frequencies and parameters, one of which is shown in Fig. 2. SNR values generally reached above 10,000, depending on the frequency and magnitude of the source activity. When the **R** covariance matrix was constructed from neighboring frequencies, SNR was high for all SSVEP frequencies and when using FWHMs above around .5 Hz (at narrower widths, the filter was simply too narrow, thereby attenuating the peak frequency). When the **R** covariance matrix was constructed from the broadband signal, SNR values were as high as when **R** was computed from surrounding frequencies, for SSVEPs above around 12-15 Hz. This is not surprising considering that the simulated SSVEP was relatively small compared to the ongoing EEG signal. Below 15 Hz, narrower filter widths increased SNR by helping to suppress endogenous EEG activity.

**Fig. 2.**
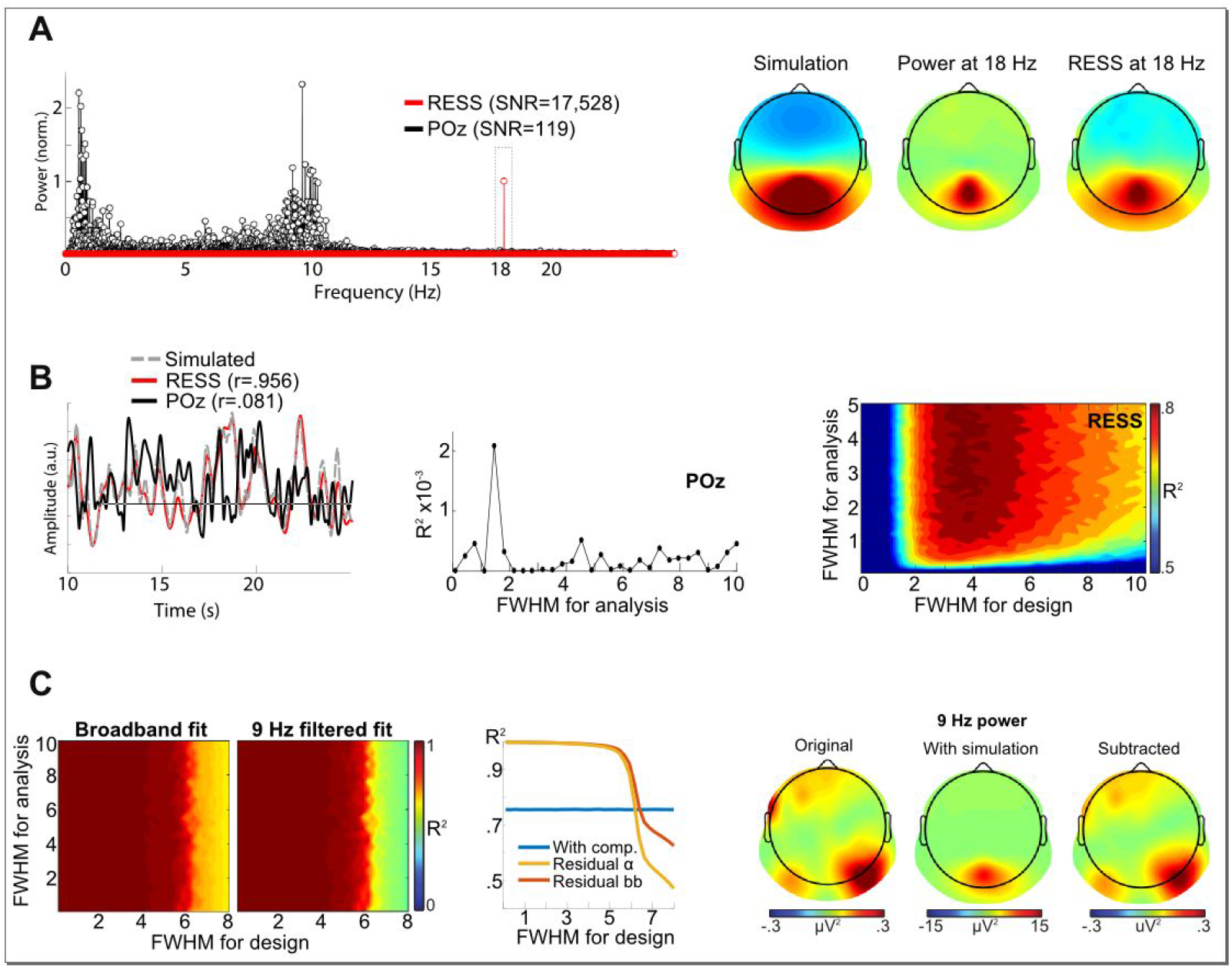
Results from simulated SSVEP. A simulated SSVEP at 18 Hz was projected into empirical resting-state EEG data. (A) The RESS method isolated the simulated signal, leading to an SNR at 18 Hz, two orders of magnitude higher than that of the “best-electrode” approach (power is normalized to 1 at the SSVEP frequency, so the interesting comparison here is the power at non-SSVEP frequencies). Topographical maps show the simulated dipole projection, the electrode power at 18 Hz, and the topographical projection of the RESS filter. (B) Time series of simulated data (dashed black line), RESS component (red line), and best-electrode (solid black line). The RESS component nearly perfectly reconstructed the simulated time series, whereas the best-electrode time series is uncorrelated with the simulated source time series. The middle and right-most plots show the effects of FWHM of the neighboring frequencies (used to design R covariance matrix), and the FWHM of the filter applied to the RESS component (after the RESS filter was applied to the raw EEG time series), on the fit between the RESS component and the simulated time series. (C) Comparison of the original data (left-most panel) and data reconstructed with the RESS component removed. A value of 1 indicates that the SSEP was perfectly removed from the data, thus a perfect separation of SSEP-related and endogenous activity. The key determinant of the accuracy of the reconstruction was the FWHM at the neighboring frequencies.

SNR computed from the best electrode increased with SSVEP frequency as the EEG signal power decreased, reaching a maximum of around 1200 for high frequencies and strong source power. Although this would normally be considered a large value for SSVEP SNR, it is an order of magnitude smaller than that obtained from RESS in the same dataset.

We next sought to determine whether the RESS and best-electrode methods could recover the time-course of the SSVEP. This is not as trivial as it initially may seem: Although the sensory stimulation may be extremely narrow-band and frequency-and amplitude-stationary, the brain's response to it has time-varying amplitude and frequency characteristics, and these non-stationarities manifest as side-lobes in the Fourier power spectrum (Fig. S1). Thus, in order to reconstruct time-varying amplitude fluctuations in the SSVEP, the filter should be wider than the stimulation frequency. But on the other hand, wide-band filtering allows non-SSVEP-related activity to contaminate the time course.

RESS performs favorably in this regard: Because the RESS filter suppresses SSEP-unrelated activity, the spatial filter can be applied to the unfiltered data, not to the narrow-band filtered data used for eigendecomposition. This allows for higher temporal precision of the RESS component. The correlation between the reconstructed SSVEP time course and the original simulated SSVEP depended on the amount of noise in the simulation, but generally was above r=.6 and was as high as r=.98 with minimal noise (Fig. 2b). The optimal filter FWHM in the simulations shown in Fig. 2 was around 4 Hz FWHM. The maximum depends in part on the spectral characteristics of the simulated SSEP magnitude, but in general we find an inverted-U-shaped relationship such that very narrow and very wide filters are suboptimal. That said, even with a very-narrow-band filter (.5 Hz FWHM) that produced overly smoothed time courses, the correlation with the simulated time course was reasonable (r > .6), and much higher than attempted reconstructions using the best-electrode approach (Fig. 2).

For time-course reconstructions, using covariance matrix R computed from the broadband signal produced slightly better results compared to using neighboring-frequencies as a reference. This is a predictable result when considering the importance of side-lobes for accurate reconstruction of the temporal non-stationarities, and that using neighboring frequencies to construct reference covariances suppresses activity at the side-lobes. Nonetheless, neighbor-frequency **R** matrices also allowed for accurate SSVEP time course reconstruction, particularly when the FWHM was relatively wide (e.g., 2-3 Hz). The point here is to avoid excessive suppression of power at immediate frequency neighbors of the stimulation frequency, because these immediate neighboring frequencies contain the temporal dynamics that are often of interest in cognitive experiments.

Simulated SSVEP time-course reconstruction using the best-electrode approach performed poorly: Correlations between time-varying power at the SSVEP frequency from the electrode that showed highest power and the simulated SSVEP time course were close to zero (Fig. 2b). The reason for the poor performance is that very-narrow-band filtering produces time courses that are too sluggish to capture the simulated temporal dynamics, while wider-band filtering fails to capture the simulated temporal dynamics due to contamination from non-SSEP neighboring frequencies.

Next, we used the RESS method to separate SSVEP from endogenous spontaneous oscillatory activity. For this, we reconstructed the EEG data using all RESS components except the one associated with the largest eigenvalue, and then compared the reconstructed data to the original EEG data (before the simulated SSVEP was added; Fig. 2c). The data with the SSVEP correlated with the original data at around r=.72, indicating the added SSVEP obscured some of the original signal. Removing the largest RESS component yielded correlations very close to 1, indicating that the SSVEP was completely removed, and the residual signal perfectly matched the original data. Of course, in real data it is difficult to confirm the accuracy of the component-subtracted residual, in part because the SSVEP may project onto more than one component. We return to this issue with the empirical data.

### Empirical EEG data

Simulations are important to demonstrate feasibility of a method, but confirmation in empirical data provides an irreplaceable examination of how a method performs under realistic conditions. We collected empirical data from one human volunteer as proof-of-principle demonstration using several different rhythmic stimulation conditions that are commonly applied in SSEP cognitive neuroscience research, or that provide difficulties for traditional SSEP data analyses.

### Visual stimulation using spatially overlapping stimuli

Centrally positioned two circles of LED lights were placed at eye level and flickered at 10 Hz and at 17 Hz (inner/outer frequencies were counter-balanced across two recordings). Inner circle LEDs were blue, outer circle LED were white. RESS successfully separated spatial patterns for 10 Hz and 17 Hz SSVEPs. Although the SSVEP at 10 Hz and 17 Hz (and their harmonics) are clear in the electrode data, RESS produced SNRs around 2-3 times larger than SNRs of the electrode data (Fig. 3a).

**Fig. 3.**
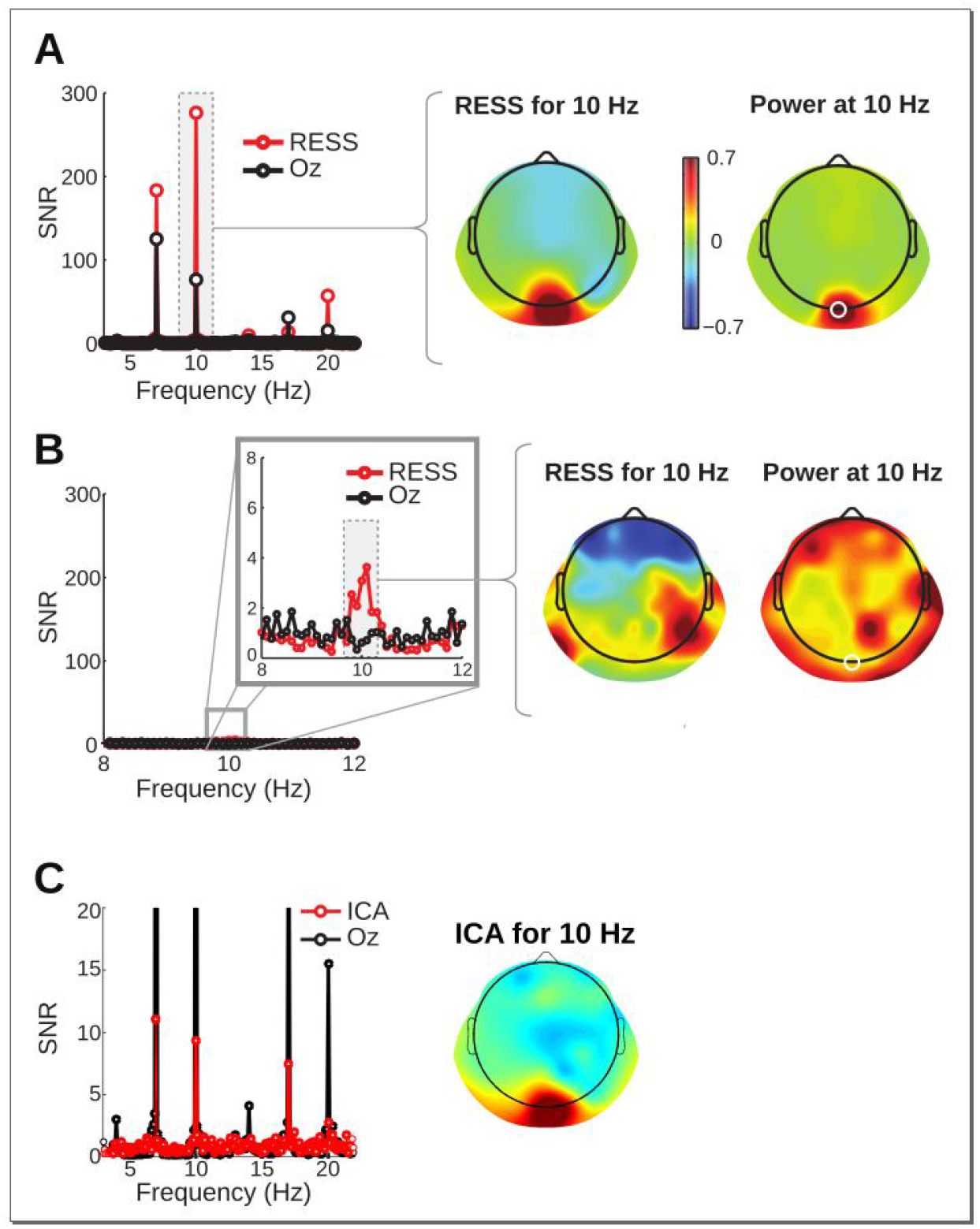
RESS applied to real data during visual flicker at 10 Hz and 17 Hz. (A) Comparison of spectral SNR and topographical maps. Note that the RESS-SNR is maximal at 10 Hz, while the best-electrode-SNR is higher at 7 Hz (this is an interference subharmonic; there was no actual sensory stimulation at 7 Hz). (B) An example of how RESS overfits data when there is no SSVEP, and how that compares to situations when real SSVEP is present in the data. The same analysis as in panel A was applied to data in which no SSVEP was present at 10 Hz. The overfitting produced a maximal SNR just under 4, compared with the real SNR of close to 300. Panel C shows that ICA on bandpass filtered data produces results that, while clearly identifying the SSVEP, had lower SNR than the best-electrode approach (electrode SNRs are cut-off to facilitate comparisons).

The increased SNR for RESS is partly a biased measure: The spatial filter is specifically designed to maximize power at the center frequency while suppressing activity at neighboring frequencies. Thus, SNRs >1 can be expected even with random data. To determine the SNR that can be expected by overfitting, we applied RESS using the same parameters as for the 10 Hz flicker, but in a condition without 10 Hz flicker (Fig. 3b). In this example, the SNR at 10 Hz was 3.6, which can be compared with an SNR of 276.3 when the 10 Hz flicker was present. This shows that although RESS may overfit data, a real effect is considerably larger.

We examined how well independent components analysis (ICA) could separate SSVEPs from two spatially overlapping flickering frequencies. The data from the 10/17 Hz experiment were narrow-band-pass filtered at flicker frequencies and subjected to an ICA decomposition using the Jade algorithm (Cardoso, 1999). In general, RESS outperformed ICA. Component selection was also less straight forward compared to RESS; for example, the largest independent component reflected blinks rather than the SSVEP response. The component that maximized power at the flicker frequency was the fourth. Fig. 3c illustrates one example of ICA compared to RESS. Comparison of RESS and ICA performance in isolating SSVEP component in other experiment conditions (not shown here) led to the same conclusion.

Flickering stimuli often produce brain responses at higher harmonic frequencies (two, three, and more times the stimulation frequency) (Kim et al., 2011), which sometimes show stronger attention modulations (Kim et al., 2007). Higher harmonic SSVEP analyses can be used to study whether different brain regions and networks might be preferentially activated by different frequencies (Heinrichs-Graham and Wilson, 2012; Lithari et al., 2016), or to study attention modulations at higher temporal precision than the base frequencies.

We applied RESS to harmonic responses (double the stimulation frequency: 20 Hz and 34 Hz) and observed strong SSVEPs (Fig. 4). Neural responses at harmonic frequencies had different topographical and anatomical distributions compared to the responses at the fundamental frequencies. At higher harmonics, SNR values could be up to seven times higher for RESS compared to the electrode data. This is important, because SSVEPs from higher harmonics often coincide with the muscle activity spectrum. Here, separating SSVEPs at harmonic frequencies using RESS was preferable: Inspection of topographical maps suggests that the electrode-level data were more sensitive to EMG noise at temporal and frontal electrodes compared to the topographical and anatomical distribution of the RESS forward model.

**Fig. 4.**
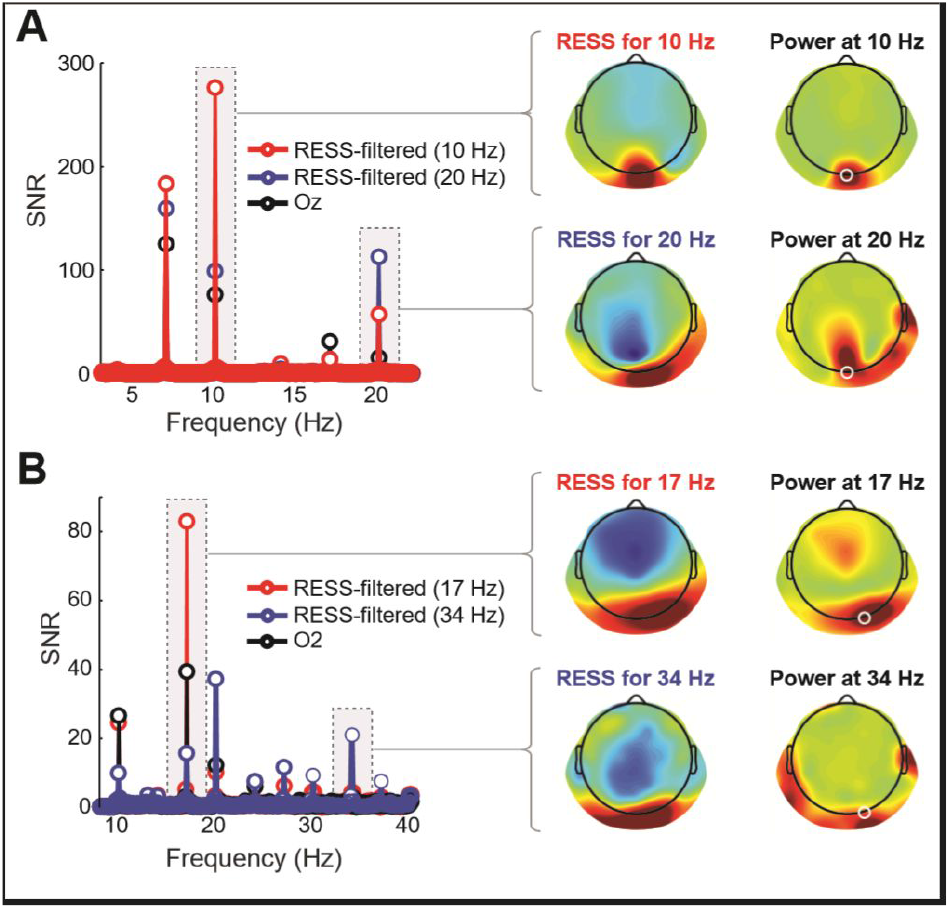
Higher harmonic SSEP analyses using RESS. Here, RESS was applied to the harmonic frequencies 20 Hz (A) and 34 Hz (B) during simultaneous 10 Hz and 17 Hz stimulation. Note the differences in topographies at the fundamental and higher harmonics.

### Visual stimulation using closely spaced frequencies

Closely spaced flicker frequencies are useful to minimize perceptual differences between different stimuli, but their corresponding SSVEP traces can be difficult to disentangle due to potential spectral leakage. To test RESS performance under these situations, we repeated the same experiment as reported above using flickering stimuli at 16/17 Hz instead of 10/17 Hz. RESS performed well, isolating different networks for the two frequencies (Fig. 5). We avoided using 17 Hz as a neighbor when computing the **R** covariance matrix for 16 Hz (and vice-versa); nonetheless, RESS produced a pair of spatial filters that doubly-dissociated the two temporal frequencies.

**Fig. 5.**
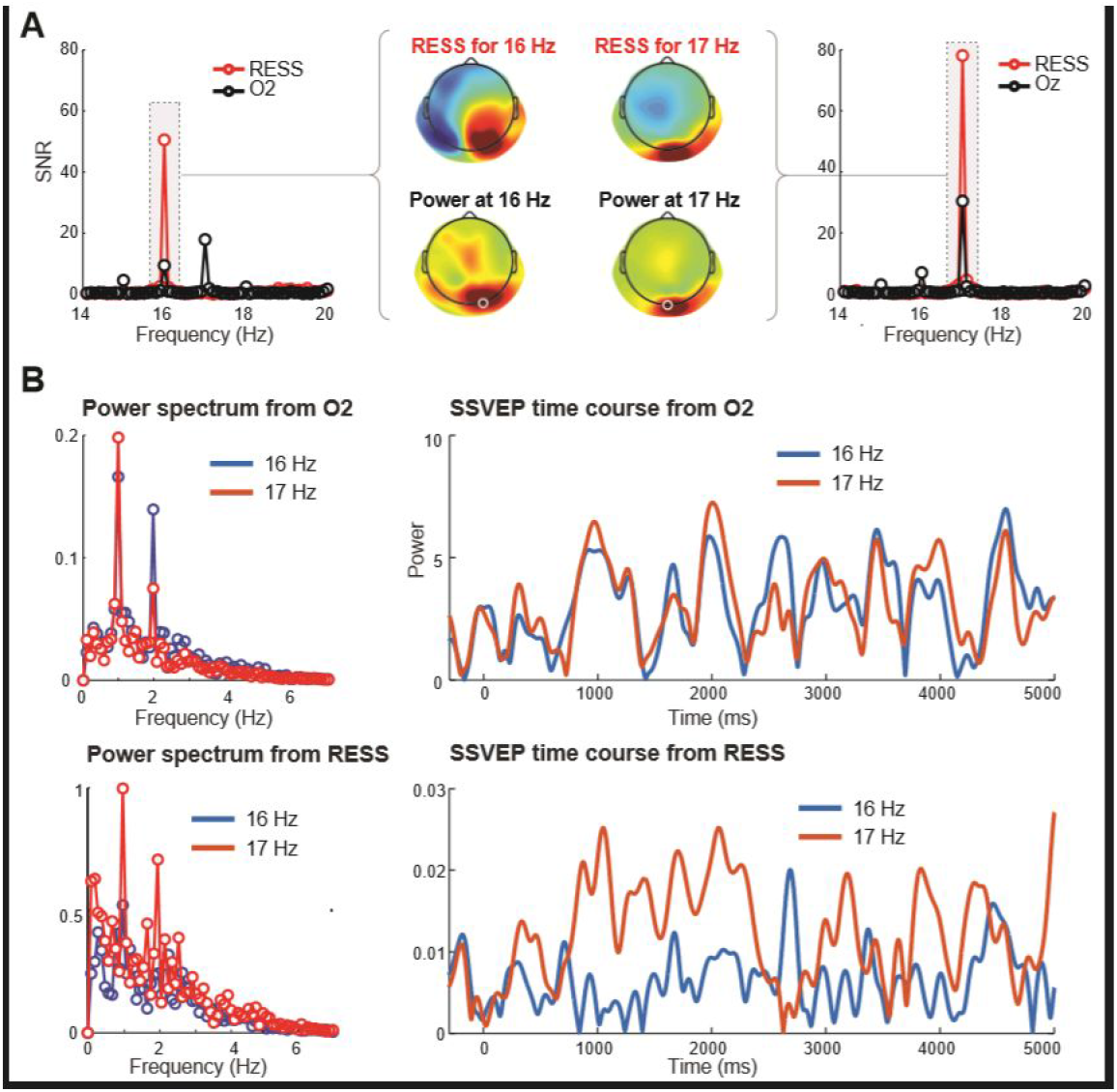
Separation of closely spaced SSVEP frequencies using RESS. (A) Power spectra and topographical maps of RESS filters during spatially overlapping simultaneous 16 Hz and 17 Hz flicker experimental condition. Note the differences in RESS topographies relative to the similarities in the electrode power topographies, and the relative suppression of power at non-SSVEP frequencies in the power spectrum of RESS component. (B) The two neighboring frequencies produced a subjective beat frequency at 1 Hz that is visible in the power spectra (left-most panels) and time-varying fluctuations in SSVEP power (right-most panels). Although the true time courses of SSVEP power fluctuations are not known a priori, the two time series were largely overlapping for the electrode data, whereas the RESS time series for 16 Hz and 17 Hz flicker were more differentiated.

When two flickering stimuli are close in frequency, they can produce a perceptual “beat rhythm” at the difference frequency, similar to interference patterns. In other words, simultaneous 16 Hz and 17 Hz flicker should produce a 1-Hz rhythm. This is observable by plotting the power time series (the envelope of the Hilbert transform) of the SSVEP time courses for the two frequencies. Both the electrode data and RESS power time courses exhibited low-frequency fluctuations and a 1-Hz power spectral peak (Fig. 5b). The electrode data power spectra were highly similar for the 16 Hz vs. 17 Hz time series, whereas RESS power time series showed clear differences between 16 Hz vs. 17 Hz. Because the SSVEP frequencies are closely spaced, any filter useful for time series examination (here, 17 Hz FWHM) cannot isolate a single SSVEP frequency, which is likely the reason why the two spectra are so similar. However, because the two RESS spatial filters were frequency-specific, a relatively wide temporal filter can be applied with minimal contamination from other, even closely spaced, flicker frequencies. This independence manifests as differences in the two power spectra (Fig. 5b).

### Flicker duration and SNR

One issue often encountered in cognitive SSEP research is the duration of the flickering stimulus. Longer durations (e.g., many seconds) ensure higher signal-to-noise ratio SSEPs, but this comes at the expense of more constrained experiment designs. Furthermore, sudden stimulus onsets produce large phasic responses that may contaminate the SSEP. We tested the ability of RESS to isolate an SSEP component using varying time windows ranging from 500 to 5000 ms, beginning at trial onset (thus including the initial stimulus-induced transient) or up to 5000 ms later (well after the stimulus-induced transient). The total stimulus duration was 10 seconds. Results are shown in Fig. 6.

**Fig. 6.**
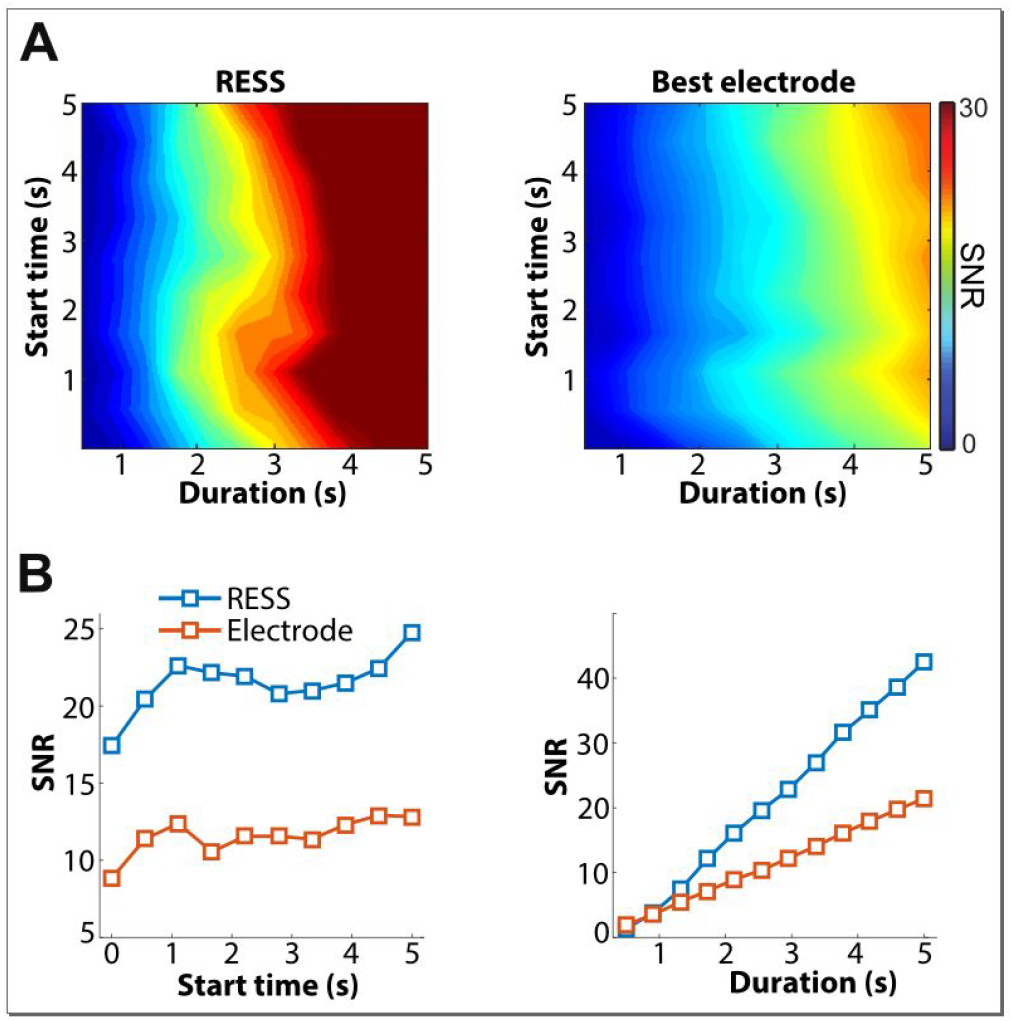
SNR by duration and start time. SNR increases when more data are used to compute power spectra, and this increase was larger for RESS compared to best-electrode. SNR also increased when the start time was later (the numbers indicate seconds after stimulus-onset), indicating that initial stimulus transients can interfere with estimation of SSVEPs. Nonetheless, SNRs were high even when including the transients in the analyses, particularly for RESS. Panel A shows the full 2D parameter space, and panel B shows averaged result over each dimension, so that RESS and best-electrode approaches can be directly compared.

Not surprisingly, SNR increased linearly with the duration of time analyzed, although this increase was steeper for RESS compared to best-electrode. SNRs were somewhat lower when including the first few hundred milliseconds post-stimulus likely due to the build-up of the SSEP response in combination with initial stimulus transients interfering with the SSVEP. Nonetheless, SNR remained high regardless of the starting time. SNRs were higher for RESS compared to best-electrode.

### Dissociating endogenous EEG from SSVEPs

The simulation results clearly demonstrated that SSVEP-related activity can be removed from the data to allow examination of non-SSVEP-related endogenous activity. It is more difficult to establish the accuracy of component subtraction in empirical data. We subtracted the 10 Hz SSVEP component from the data and found that the stimulus onset and offset responses were preserved, while the increased power during the long stimulus presentation was strongly attenuated (Fig. 7). Topographical maps showed that removing the largest RESS component affected only a small region of the topography. This shows that 10 Hz activity was not simply globally suppressed; instead, the 10 Hz activity that was removed was both topographically and temporally localized.

**Fig 7.**
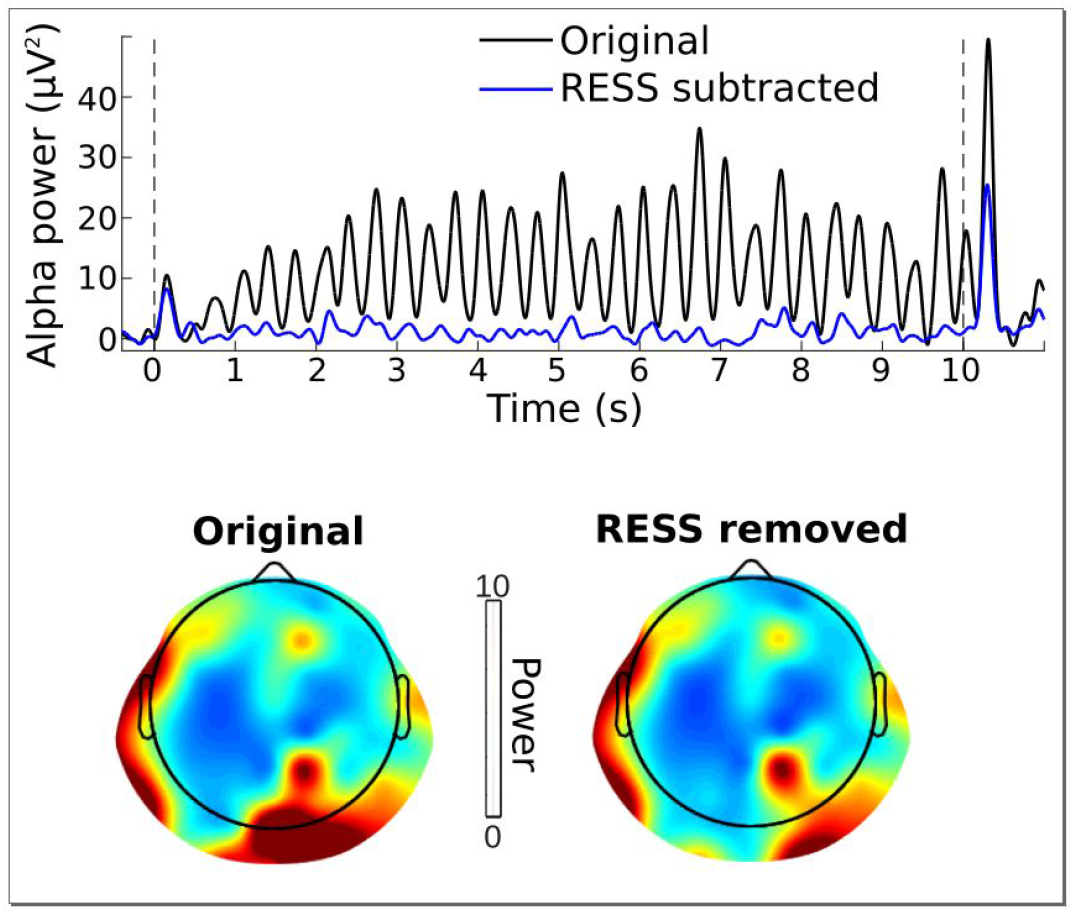
Separation of SSVEPs and endogenous EEG using RESS. The EEG data from 10 Hz visual flicker stimulation showed a strong SSVEP modulation (black line, taken from electrode Oz). Identifying and subtracting the largest RESS component yielded an alpha power time course (blue line) that was comparable in initial stimulus onset and offset to the original data, and which was topographically similar except in central posterior sites. The goal here was to “remove the SSVEP” from the EEG data, although this result must be interpreted with caution. See Fig. 3a for a topographical map of the RESS component that was removed.

### Noisy electrodes

EEG recordings occasionally have noisy or bad electrodes. For single-electrode-based analyses, bad electrodes can be simply ignored or interpolated. But because RESS is a weighted combination of all electrodes, leaving bad or excessively noisy electrodes in the dataset might have a negative impact.

We investigated the effects of electrode noise by replacing the EEG activity at electrodes Cz, Pz, and Iz with large-amplitude random white noise (mean 0, standard deviation 1000). The key comparison is the SNR of the SSVEP with the noisy electrodes in the data vs. removing the noisy electrodes. Here we focus on the 17 Hz SSVEP, but the findings are comparable for other frequencies and conditions. The topography of 17 Hz power was dominated by the noise electrodes, although of course the SNR spectrum at Oz was unchanged. The RESS forward model was cleaner when removing the electrodes (Fig. 8), and SNR at 17 Hz increased from 81.36 (with noisy electrodes) to 83.28 (without the electrodes). Thus, in this case, removing the noisy electrodes improved SNR by 2.3%.

**Fig. 8.**
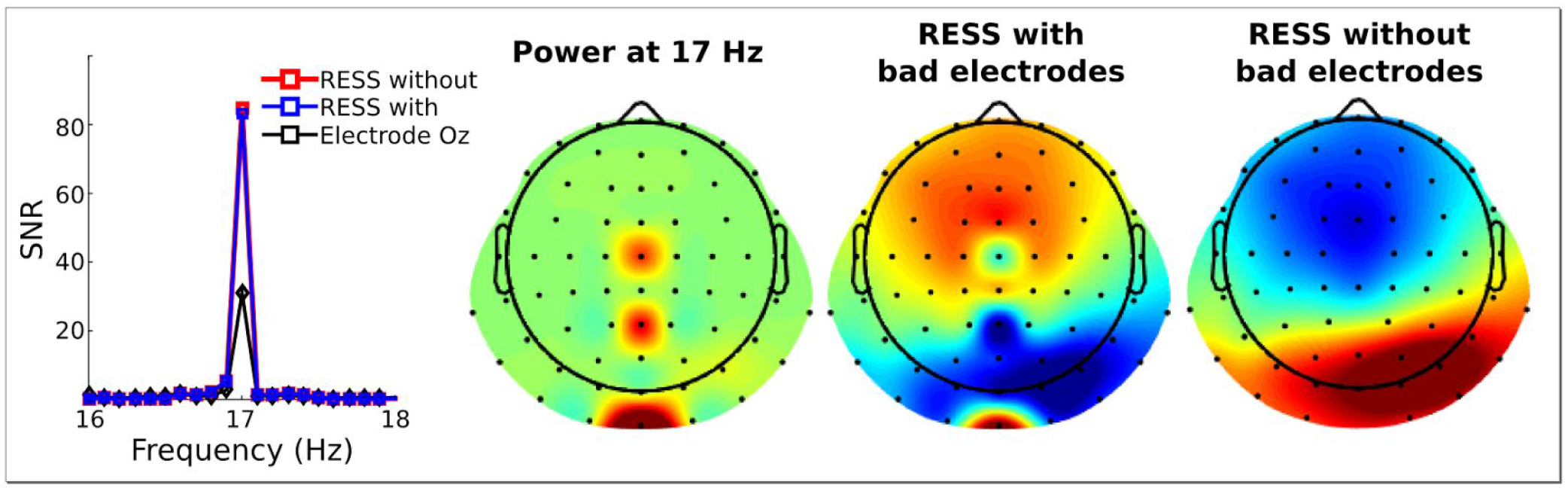
Effect of noisy electrodes on RESS components. Replacing three electrodes (Cz, Pz, Iz) with large-amplitude noise affected topography of the RESS component, but had minimal deleterious effects on the SNR. The right-most topographical map and red SNR trace shows results after removing the three noisy electrodes from the data.

We tested effects of other levels of noise, summing noise on top of (instead of replacing) the EEG data, and noisy electrodes in other datasets (not shown here). Our observations across these tests leads to a consistent conclusion: Removing noisy electrodes from the dataset improves SNR slightly, while the spatial projections become cleaner and more easily visually interpretable.

### Anatomical localization

The forward model of the RESS spatial filters can be projected onto the brain if one has a suitable forward model that would be used for, e.g., beamforming or minimum-norm estimation. Fig. 9 illustrates a few examples of scalp and brain projections. Although localization of EEG generators contains several sources of uncertainty, such anatomical maps often provide useful information, as well as visually compelling results.

**Fig. 9.**
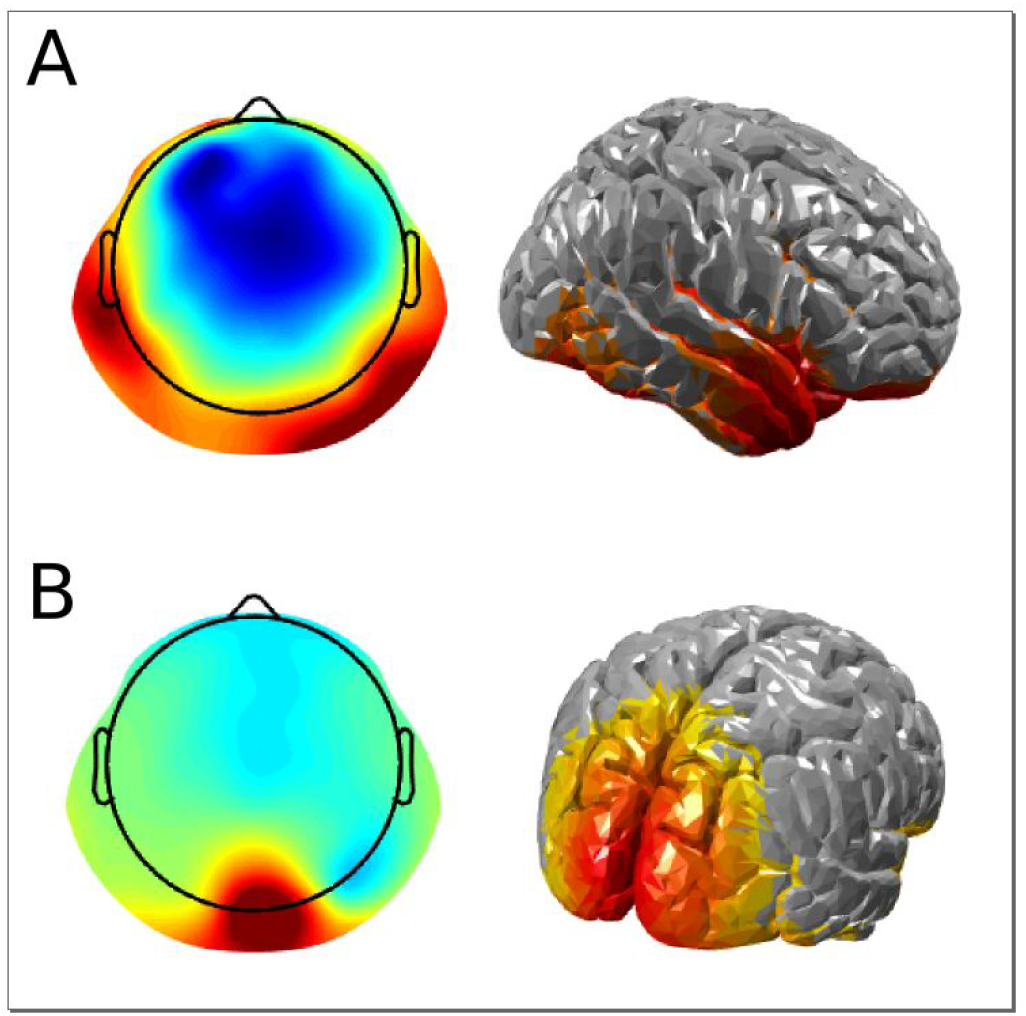
Putative anatomical distributions can be computed from RESS components. (A) RESS component taken from 40 Hz binaural auditory stimulation. (B) RESS component taken from 10 Hz visual stimulation (same component as is shown in Fig. 3a).

## Discussion

In the course of analyzing our SSVEP data and discussing SSVEP analyses with other researchers, we identified the six analysis challenges described in the Introduction and sought analysis techniques that could address these challenges. We found that no single technique provided satisfactory results, but that a blend and extension of several existing techniques performed very well. The RESS method outlined here combines several existing techniques and is firmly grounded in decades of development in signal processing and multivariate source separation methods (Parra and Sajda, 2003; Särelä and Valpola, 2005). It most closely follows the work of Nikulin et al. (Nikulin et al., 2011), Cheveigne et al. (de Cheveigné and Arzounian, 2015), and Dmochowski et al., (Dmochowski et al., 2015). The contribution of the present work is the optimization of generalized eigendecomposition to the SSEP-related analysis issues identified in the Introduction.

The general idea of RESS is simple: Find a vector that maximally differentiates two covariance matrices—one corresponding to the to-be-maximized signal and one corresponding to the to-be-minimized reference—and then use that vector as weights to combine the data from all electrodes. The subtleties of the method come from selecting appropriate parameters, including the temporal filter bandwidths, frequencies to use as reference, and methods of preprocessing the data. For this reason, we encourage using simulations and pilot data to find suitable parameters given the sensory stimulation frequencies, expected topographical maps, etc. We stress that RESS should not be used as an automated (“blind”) procedure that is applied without consideration. Instead, one must approach the data carefully and optimize the parameters based on the goals of the analyses, details of the experiment design, and assumptions about the underlying neurocognitive temporal dynamics.

Simulations and proof-of-principle empirical data clearly demonstrate the advantages of RESS over the best-electrode picking procedure prevailing in the SSEP literature. Indeed, even when RESS provides only marginal improvement over the best-electrode approach in terms of SNR (this can happen with large stimuli and long presentation times), RESS still avoids the issue of having to select specific electrodes, which may differ by individual, flicker frequency, and experiment condition. Although empirical SSVEP data used here was obtained using arrays of LEDs which elicited a large-amplitude SSVEPs, we have also applied RESS to datasets in which smaller and shorter duration stimuli were presented on standard (60-100 Hz refresh) monitors, and obtained satisfying results. The main potential danger of RESS is overfitting noise, which we discuss in the next section.

### Overfitting: Uses and missuses

There is a danger of overfitting when using this method—or indeed, when using any form of “guided” filtering technique. Essentially, we are using RESS to search through an N-dimensional space (where N is the number of electrodes) for a vector that maximizes what we explicitly designed the filter to maximize. This method applied to pure noise would still produce a bump in the power spectrum, which was illustrated in Fig. 3b.

There are at least two appropriate ways to use RESS (and related methods) to optimize data analysis while avoiding overfitting. The first way is to define the spatial filter based on the data from all experiment conditions and then apply the spatial filter separately to each condition. For example, imagine that an experiment has conditions attend-flicker and ignore-flicker (both at the same frequency). The general presence of a RESS-defined SSVEP effect should be interpreted with caution because such an effect could be due to overfitting noise, but differences between attend and ignore conditions are safely interpretable. A second way to avoid overfitting would be to define the spatial filter based on independent data, either from a separate task or at least from different trials in the main task.

### Related to overfitting

It is possible to develop an automatic search algorithm to find a set of parameters that maximizes SNR. For example, a script could loop over different ranges of FWHM and neighbor distances and pick the parameter set that produces the largest SNR. However, here one must be particularly mindful of overfitting. The best way to select analysis parameters is a combination of simulations, testing in empirical pilot data, and common sense. Once a set of parameters is selected, it should be applied to all experiment conditions (although different frequencies may require different parameters), individuals, groups, etc.

On the other hand, overfitting noise can be used advantageously, as a method of statistical comparison. For example, computing the RESS power spectrum on hundreds or thousands of data time series from the same experiment but with sensory flicker at different frequencies can produce a comparison distribution against which the true RESS power spectrum (from trials at which the flicker frequency matches the RESS filter frequency) can be compared. This is comparable to null-hypothesis testing.

### Recommendations for SSEP experiment design and analysis when using RESS

RESS and other spatial filters work best when the sources have spatially distinct projections. There is evidence in the literature that, at least in the visual domain, different stimulus frequencies activate different networks (or at least, produce different topographies; e.g. Heinrichs-Graham & Wilson, 2012; Lithari et al., 2016). It is also known that some low-level visual features, such as different phases of Gabor patches, produce different topographies as evidenced by topographical decoding accuracy (Wardle et al., 2016). Having spatially segregated stimuli is perhaps the best way to facilitate different topographies (Vanegas et al., 2013), which in turn will maximize the efficacy of the spatial filter.

The time series data must have the correct number of points (zero-padding when necessary) to extract the precise stimulation frequency from the data. For example, if the stimulus flickers at 15.4 Hz but the FFT is computed with 1 Hz resolution, the neural response to the flicker will be suboptimally reconstructed.

When possible, define **S** and **R** covariance matrices based on data from the same time periods. This will hold constant all cognitive and task-related dynamics, allowing the spatial filter to maximize frequency differences. For example, if the **S** covariance matrix is computed from a demanding cognitive task while the **R** covariance matrix is computed from a task-free resting period, then the filter will maximize both frequency and task-vs.-rest differences, and will therefore be less specific.

Similarly, it is better to define the **R** matrix from stimulation-neighboring frequencies, because this will help suppress endogenous activity in that frequency band. For example, muscle artifacts will contaminate a 25-Hz SSEP, but because EMG noise has a relatively broad frequency spectrum, computing the **R** matrix from 23 Hz and 27 Hz will effectively suppress EMG activity. On the other hand, if time course analyses will be performed, neighboring frequencies should not be too strongly suppressed. In this case, we recommend using wider FWHM at neighboring frequencies compared to the peak frequency (e.g., .5 Hz for the peak and 2 Hz for the neighbors).

For the spatial filter to remain valid, the spatial location and the temporal frequency of the entraining stimulus must be constant. RESS is not suitable, for example, if a flickering visual is moving around the monitor, not is it suitable for frequency chirps (time-varying frequency changes).

Concerning data cleaning and preprocessing, we recommend removing trials or sections of data that contain excessive artifacts, but not subtracting independent components. In our experience, removing artifactual independent components led to comparable or worse performance of RESS, and reduced the rank of the covariance matrices, which may reduce the accuracy of the eigendecomposition. Bad or excessively noisy electrodes should not be interpolated, because this produces reduced-rank covariance matrices. Instead, removing excessively noisy electrodes prior computing RESS is advisable. Furthermore, it is not necessary to have the same number of electrodes in each subject because the electrodes themselves are not considered in data analyses, only the components. That said, the number of electrodes should be held roughly constant over subjects, because the number of electrodes in part defines the quality of the decomposition. Excessively noisy electrodes are generally suppressed by RESS, so it may not be necessary to remove them, also considering that SNR seems to increase only slightly (around 0-3% in our tests). On the other hand, if differences between experiment conditions are subtle, even minor additional improvements in SNR may beneficial. If putative anatomical localization is an important consideration in the interpretation, then removing excessively noisy electrodes is recommended.

### Other general recommendations

The quality of the RESS method hinges on the quality of the covariance matrices. A high-quality covariance matrix comes from clean data with many time points. Use as much data (that is, as many time points) for the covariance matrix as possible, while using only data that include steady-state stimulation. We have encountered poor performance (e.g., complex eigenvectors or close-to-singular matrices) with single-precision data, thus making sure that the data are in the highest computer precision possible is important. If complex eigenvectors are obtained, the neighboring frequencies that make the **R** matrix can be moved a bit further away and/or their FWHM can be made broader.

In general, it is ideal to have a single set of parameters for all frequencies. However, we have observed that over a large range of frequencies (e.g., 3 Hz to 80 Hz), a single set of filtering parameters is unlikely to be suitable. For example, **R** matrix frequencies might need to be further apart from the SSEP frequency at lower frequencies and closer to the SSEP frequency at higher frequencies. Piloting with a subset of the data will help determine appropriate parameter ranges. However, it is important that parameters (and the spatial filters) are matched across experiment conditions to prevent biases.

### Relationship to other spatial filters

We found that RESS outperformed ICA for isolating SSEP components (see also Nikulin et al., 2011). However, it should be cautioned that the superiority of RESS over ICA is trivial. “Guided” source separation techniques—particularly those that involve carefully selected maximization **(S** matrix) and minimization **(R** matrix) goals will generally always outperform blind source separation techniques that have the goal of decomposing the entire signal. One can think of ICA as *fitting* the data, and of RESS and other source separation techniques as *overfitting* the data. Despite this caution, the comparison with ICA demonstrates that the brain produces spatiotemporal dynamics in response to flickering stimuli, and that commonly used EEG data analysis techniques such as single-electrode or ICA may be poorly suited to uncover these dynamics.

Our method is partly inspired by the spatiospectral decomposition (SSD) method (Nikulin et al., 2011). Two advantages of RESS are that the temporal bandpass filtering is narrower and that the spatial filter is applied to the broadband data. These features increase spectral specificity while allowing for more flexibility in reconstructing the temporal fluctuations of the RESS component. In general, there are no one-size-fits-all data analysis methods, and exceptional cases (like SSEPs) often require specific procedures.

We also tested a related technique referred to as joint decorrelation (JD; de Cheveigné and Parra, 2014). The idea of joint decorrelation is to sphere the data by normalizing the covariances, and then apply a “temporal bias filter” to identify patterns of covariance that match the filter. In our case, the filter was a sine wave at the stimulation frequency. JD produced acceptable results that were often comparable with RESS (results not shown). However, JD has two disadvantages with respect to SSEP. First, the temporal bias filter is a stationary sinusoid, whereas the brain's response to a rhythmic stimulus is neither perfectly stationary nor is it perfectly sinusoidal. Second, JD is not designed to suppress power at neighboring frequencies, meaning it captures a mixture of SSEP and endogenous activity at the same frequency. Again, this is not a criticism of JD in general; instead, it reflects that a specific experimental method like SSEP requires a correspondingly specific data analysis technique.

There are myriad other spatial filters that have been and could be applied to SSEP data. Other spatial filters, for example, are based on maximizing the difference between trial-averaged and single-trial covariances (Dmochowski et al., 2015; Kuś et al., 2013). This approach, however, assumes that (1) the timing of the flicker with respect to trial onset is the same on every trial (in other words, the exogenous sensory stimulus must be “phase-locked”), and (2) the single-trial variance reflects noise to be minimized. These two assumptions may be held in some situations, but they force stringent constraints that limit experiment design and analysis possibilities that make them less useful in many cognitive neuroscience applications, particularly with regard to within-subject or single-trial analyses. For example, imagine an experiment in which the background flickers while a visual stimulus can appear with random temporal jitter. In this case, the sensory flicker is “non-phase-locked” with respect to stimulus onset, and any spatial filter designed to maximize the trial-averaged SSVEP would perform poorly.

The Laplacian is often used to increase signal-to-noise characteristics and increase topographical specificity by attenuating volume conducted activity (Cohen, 2014; Kayser and Tenke, 2015; Winter et al., 2007). The Laplacian is also applied to SSVEP data with success (Srinivasan et al., 2006). However, given that Laplacian increases topographical specificity, topographical differences across different flicker frequencies and individuals may become more pronounced. This, in turn, makes careful electrode selection more important if “best-electrode” approach is used. We have tested RESS on several datasets with and without applying the Laplacian (results not shown) and found no noteworthy differences in SNR.

### Interpreting RESS components

RESS is simply a linear weighted combination of electrodes, where the weights are defined according to maximizing the distance between two covariance matrices. There are no constraints regarding spatial smoothness or anatomical localization. Although this is advantageous because it makes the filter simple, fast, and robust, it also means that one cannot simply assume that each component corresponds to a single dipolar neural generator (although this is possible).

On the other hand, unlike principle components analysis, in which the vectors are selected to be all pairwise orthogonal, RESS vectors are not orthogonal. This is because although **S** and **R** matrices are both symmetric, **R-1S** is not symmetric (orthogonal basis vectors are guaranteed only in eigendecomposition of symmetric matrices). This means that RESS is conceptually closer to demixing (like ICA) as opposed to decorrelating (like PCA); multiple correlated neural generators are not forced into orthogonal statistical components.

We have observed RESS to provide (in different frequencies and datasets) both spatially very localized anatomical estimates, and spatially distributed estimates. Thus, we recommend interpreting RESS results more in terms of statistical or signal-space components rather than in terms of single anatomically isolated neural generators. In other words, there is a subspace “pocket” inside the global data space that contains the SSEP-related dynamics, and the purpose of RESS is to point the researcher in the direction of that pocket (see Cheveinge and Parra, 2005, for more discussion of this point). In more neurobiological terms, each RESS component can be thought of as capturing a functionally cohesive—possibly spatially restricted or possibly spatially distributed—brain network that responds to rhythmic sensory stimulation in a statistically similar way.

## Acknowledgements

MXC is funded by an ERC-StG 638589

